# Super-resolution imaging reveals changes in *Escherichia coli* SSB localization in response to DNA damage

**DOI:** 10.1101/664532

**Authors:** Tianyu Zhao, Yan Liu, Zilin Wang, Rongyan He, Jia Xiang Zhang, Feng Xu, Ming Lei, Michael B. Deci, Juliane Nguyen, Piero R. Bianco

**Affiliations:** Xi’an Institute of Optics and Precision Mechanics, Chinese Academy of Sciences, State Key Laboratory of Transient Optics and Photonics, Xi’an, 710119, China; The Key Laboratory of Biomedical Information Engineering of Ministry of Education, School of Life Science and Technology, Xi’an Jiaotong University, Xi’an, Shaanxi 710049, China; Bioinspired Engineering and Biomechanics Center (BEBC), Xi’an Jiaotong University, - Xi’an, Shaanxi 710049, China; Center for Single Molecule Biophysics and, Department of Microbiology and Immunology, University at Buffalo, Buffalo, NY, USA; Department of Pharmaceutical Sciences, School of Pharmacy, University at Buffalo, Buffalo, NY, USA

**Keywords:** single strand DNA binding protein, *E. coli*, DNA repair, DNA replication, super-resolution microscopy, SSB interactome, OB-fold

## Abstract

The *E. coli* single stranded DNA binding protein (SSB) is essential to viability. It plays key roles in DNA metabolism where it binds to nascent single strands of DNA and to target proteins known as the SSB interactome. There are >2,000 tetramers of SSB per cell with perhaps 100-150 associated with genome at any one time, either at DNA replication forks or at sites of DNA repair. The remaining 1,900 tetramers could constantly diffuse throughout the cytosol or be associated with the inner membrane as observed for other DNA metabolic enzymes such as DnaA and RecA. To visualize SSB directly and to ascertain spatiotemporal changes in tetramer localization in response to DNA damage, SSB-GFP chimeras were visualized using a novel, super-resolution microscope optimized for visualization of prokaryotic cells. Results show that in the absence of DNA damage, SSB localizes to a small number of foci and the excess protein is observed associated with the inner membrane where it binds to the major phospholipids. Within five minutes following DNA damage, the vast majority of SSB disengages from the membrane and is found almost exclusively in the cell interior. Here, it is observed in a large number of foci, in discreet structures or, in diffuse form spread over the genome, thereby enabling repair events. In the process, it may also deliver interactome partners such as RecG or PriA to sites where their repair functions are required.

## Introduction

The *Escherichia coli* single stranded DNA binding protein (SSB) is an essential protein that plays a central role in DNA metabolism (1–3). It binds to single-stranded DNA (ssDNA) non-specifically and with high affinity (4, 5). SSB also interacts with an array of at least fourteen proteins that has been termed the SSB-interactome (6, 7). These two seemingly disparate roles are intimately linked via the linker domain of the protein as explained below (8).

SSB exists as a stable homo-tetramer (9). Each 178 amino acid length monomer can be divided into two domains defined by proteolytic cleavage: an N-terminal portion comprising the first 115 residues and a C-terminal domain that includes residues 116 to 178 (4). The well conserved N-terminal or core domain contains elements critical to tetramer formation and the oligonucleotide/oligosaccharide binding-fold (OB-fold) critical to the binding of ssDNA (4, 10). Importantly, the OB-fold is structurally similar to eukaryotic Src homology 3 (SH3) domains (11). These domains are ~50 residue modules that are ubiquitous in biological systems and are well known for their ability to bind PXXP motifs present in interacting partners (12–15).

The C-terminal domain of SSB can be further subdivided into two sub-domains: a sequence of approximately 54 residues comprising residues 116 to 170, that is known as the linker and, the last 8 residues which are known as the acidic tip (3, 16). The linker contains several structural elements critical to its function, including PXXP motifs (8, 17, 18). Critical to SSB function, the PXXP motifs in the linkers of one tetramer bind to the OB-fold(s) present in a nearby tetramer thereby facilitating cooperative ssDNA binding (8, 17, 18). Furthermore, as OB-folds have been identified in each SSB interactome partner, a similar binding mechanism may be employed to mediate SSB-partner interactions critical to genome stability.

In exponentially growing cells, there are more than 8,000 SSB monomers per cell (19). In contrast, the levels of interactome partners are significantly lower with for example, as few as 7-15 RecG molecules per cell being observed. In these same cells there are on average 2-4 DNA replication forks per cell. At each fork, there is 0.5 to 1kb of ssDNA available (20). Using a site size of 40 nucleotides occluded per tetramer, there would be on average 25 tetramers bound per fork, or 100 per cell. This means that at any one time there are as many as 1,900 free SSB tetramers. These may be freely diffusing in the cytoplasm or associated with the inner membrane in a “storage form” as suggested previously (7).

To determine whether SSB localizes to the inner membrane, a novel super-resolution microscope was developed to enable clear visualization of SSB-GFP localization. Results show that in the absence of exogenous DNA damage, SSB localizes to the inner membrane and to foci (likely sites of DNA replication). *In vitro* binding studies show that SSB binds preferentially to liposomes comprised of the dominant phospholipids of the *E. coli* inner membrane. Consequently, we propose that membrane association occurs via binding to these phospholipids *in vivo*. Following DNA damage, SSB rapidly disengages from the membrane, with the majority of the GFP signal being associated with the genome. Furthermore, once damage has been repaired, SSB returns to the inner membrane. These findings suggest a control mechanism for maintaining SSB at the inner membrane utilizing phospholipid-OB-fold interactions and a means for SSB to rapidly deliver repair enzymes to the genome when their actions are required.

## Materials and Methods

### Materials

Coverslips (No. 1.5; 22×22mm) were purchased form VWR and used without further cleaning. All chemicals were reagent grade, made up in Nanopure water and passed through 0.2μm pore size filters. LB and M63 media were made according to (21). As experiments were done both in China and the USA, the sources of chemicals were different. Therefore, these are listed according to location.

(A) China. Thiamine Hydrochloride was from Solarbio (Beijing, China). (NH_4_)_2_SO_4_, KH_2_PO_4_, FeSO_4_·7H_2_O and KOH were from Zhiyuan Chemical Reagents (Tianjin, China). Casamino acids were from Sangon (Shanghai, China). D-(+)-Glucose was from MP (USA). D-lactose monohydrate was from Macklin (USA). Mitomycin C was from Roche (USA). Poly-L-lysine was from Sigma (USA) and FM4-64 was from Invitrogen (USA).

(B). USA. The λDE3 lysogenization kit was from Novagen. Yeast Extract and tryptone were from Becton Dickinson and Company (MD, USA). NaCl and lactose were from J.T. Baker (NJ, USA). Ampicillin was from Fisher (NJ, USA). IPTG was from OmniPur (NJ, USA). Kanamycin and chloramphenicol were from Sigma (MO, USA). Glucose was from Mallinckrodt (KY, USA). Primestar Max DNA polymerase and In-Fusion^®^ HD Cloning Plus was from Clontech (CA, USA). Plasmid purification and gel purification kits were form Omega-Bio tek (GA, USA). Poly-L-lysine was from either Sigma or Mallinckrodt. Lipids were purchased from Avanti Polar Lipids (Alabaster, AL). Lipids in this study included 1,2·-di-(9Z-octadecenoyl)-*sn*-glycero-d-phospho-(1’-*rac*-glycerol) (DOPG), 1,2-di-(9Z·-octadecenoyl)-*sn*-glycero-d-phosphocoline (DOPC), 1,2-di-(9Z-octadecenoyl)-*sn*·-glycero-3-phosphoethanolamine (DOPE) and 1,1’,2,2’-tetra-(9Z-octadecenoyl) cardiolipin (CL).

### Methods

#### Recombineering

Bacterial strains used are listed in the Supplementary Information, Table I. AG1/pCA24N-his-SSB-GFP was provided by the National Bioresource Project of National Institute of Genetics, Japan (22). MG1655/pKD46 was purchased from the Coli Genetics Stock Center at Yale University (23). pKD46 contains the lambda *red* and *gam* genes under the control of the arabinose promoter. The parent strain for recombineering, MG1655 *ΔlacY::Kan*, was transformed with pKD46 and plated at 30°C. A single colony was selected and grown overnight at 30° in LB + glucose (0.2% final) and ampicillin (250μg/ml). The next day, the culture was diluted 1:100 into LB containing ampicillin and arabinose (1mM final), grown to an OD_600_=0.4 and processed according to (24).

**Table I.** SSB binds to liposomes

| Liposome type<sup>a</sup> | $K_d$ (μM)<sup>b</sup> |
| --- | --- |
| 100% PG | $0.78 \pm 0.25$ |
| 80:20 PG:CL | $0.48 \pm 0.11$ |
| 100% PC | >30 |
| 90:10 PC:CL | $0.44 \pm 0.11$ |
| 70:30 PC:CL | $0.35 \pm 0.10$ |
| 80:20 PC:PG | $0.79 \pm 0.21$ |
| 80:20 PC:PE | $0.56 \pm 0.20$ |

- Liposomes were constructed as described in the Materials and Methods. The ratios indicate the percentage of each phospholipid present in the liposomes being tested. PG, phosphatidylglycerol; CL, cardiolipin; PC, phosphatidylcholine; PE, phosphatidylethanolamine. - Binding of liposomes to SSB was done by monitoring changes in the intrinsic fluorescence of the protein that occurs on binding. Data represent several experiments done on different days. Analysis was done as described previously (27).

The plasmid pCA24N-his-SSB-GFP was amplified in PCR using primers PB682 and 683 for insertion into *flgA* or PB684 and 685 for insertion into *flgE* (sequences provided in Supplementary Information). Following the first round of PCR, the fragments were gel purified. The final products were ethanol precipitated, resuspended in TE buffer and electroporated into cells using an Eppendorf Electroporator 2510 in 1mm electroporation cuvettes (Bio-Rad).

Chloramphenicol resistant colonies were grown overnight at 37°C in LB containing antibiotic (25μg/ml) and glucose (0.2% final). The next day, cultures were diluted 100-fold into fresh LB containing chloramphenicol only, grown at 37°C for 3 hours and IPTG added to 1mM final to induce expression of the SSB-GFP fusion and grown for an additional 3 hours. Cells were subjected to centrifugation at 5,000xg for 5 minutes, resuspended in 1/10^th^ volume of 10mM MgSO_4_ and GFP fluorescence assessed using a Cary Eclipse fluorimeter (Ex488/Em509).

#### Cell growth for fluorescence microscopy

Cell growth was done at 37°C with vigorous shaking. Overnight cultures were grown separately in either M63 containing 0.4% glucose and casamino acids or LB containing 0.4% glucose. The next day, cells were diluted 1:100 into fresh media of the same type, grown for 3 hours and processed for imaging as described previously (25) or treated with MMC and FM4-64 (see below).

#### Treatment with mitomycin c

Tubes of MMC were thawed just before use and added to cultures as follows. Following growth for 3 hours, 5 mL cultures were divided into 4 tubes containing 1mL of bacteria each. Tubes were treated separately with either none; 0.2, 0.4, or 1.2μg/ml MMC (final), for 30 minutes. Cells were then subjected to centrifugation and processed for imaging.

*To visualize the inner membrane*, FM4-64 was added to a final concentration of 2μM and allowed to bind for 15 minutes. When cells were treated with MMC, the membrane dye was added 15 minutes after the DNA damaging agent, followed by an additional growth of 15 minutes (total MMC exposure – 30 minutes; total dye time −15 minutes). Thereafter cells were subjected to centrifugation, deposited on coverslips and imaged.

#### Image analysis

Details of the image reconstruction procedure are provided in the Supplementary Information. The resulting images were background subtracted only and not enhanced by deconvolution. These were then analyzed using Image-Pro Premiere v9.3 (Media Cybernetics) and Fiji (26).

#### Liposomes and liposome binding

Lipids or lipid mixture were dissolved in chloroform, dried down to a thin film and placed under high vacuum to remove traces of solvent. The lipid films (2 μmol) were re-hydrated with 1 mL of R buffer (20 mM Tris, 0.1 mM EDTA, pH 7.4) at 60°C and vortexed. The liposome suspensions were sonicated (Branson 1800 bath sonicator) at 60°C for 20 min. The liposomes were extruded (Avestin LiposoFast Extruder) through a 100 nm polycarbonate membrane (Avanti Polar Lipids) at 60°C. The following compositions of liposomes (expressed as molar ratios) were used in this study: 100% DOPC, 100% DOPG, 95:5% DOPC:CL, 90:10% DOPC:CL, 80:20% DOPC:CL, 70:30% DOPC:CL, 80:20% DOPC:DOPE, 80:20% DOPC:DOPG and 80:20% DOPG:CL.

Binding of liposomes to SSB was determined by monitoring the intrinsic fluorescence of the protein at 350nm in a Cary Eclipse spectrofluorometer (Varian). Reactions were done at 37°C in a volume of 500μl. Solutions were continuously stirred and contained 10mM Tris-HCl (pH8.0), 1mM DTT, 10mM NaCl and 1μM his-SSB protein (monomer). The intrinsic fluorescence of SSB was monitored using an excitation wavelength of 290nm and then scanning the emitted fluorescence from 300-400nm, with a maximum being observed at 350nm. At various time intervals, aliquots of liposomes were added, mixed thoroughly using a pipettor and incubated for 8 minutes with stirring prior to scanning the sample for fluorescence changes. If binding occurred, the intrinsic fluorescence of SSB decreased, similar to that observed for ssDNA binding (25). An 8-minute time point was determined experimentally as this was the minimum time required to observe the maximum fluorescence decrease. When lipid dilutions were required these were done using reaction buffer and were made immediately prior to use, and discarded once the experiment was complete. As the Cary Eclipse has a multicell transport, two SSB identical reactions were done in adjacent cuvettes simultaneously with a third containing buffer only. The change in fluorescence in this third sample was used to account for light scattering changes that occurred in response to each addition by subtracting values of fluorescence that we measured for liposome-only titrations into buffer. In addition, the resulting fluorescence signals measured following each liposome addition were corrected for dilution of SSB. The resulting data were analyzed exactly as described previously (27). Corrected data were fit to one-site specific binding equation: Y= (B_max_ × X)/(K_d_+X·). Curve fitting was done using Prism v6 (GraphPad software).

## Results

### A novel super-resolution microscope for low light imaging

To visualize autofluorescent-tagged proteins in *E. coli* under conditions of low excitation intensity (≤1W/cm^2^), a novel super-resolution imaging system was developed (see Supplementary Information Figures 1 and 2). This system is comprised of a synchronized, structured illumination microscope (SIM) and a recently developed image reconstruction algorithm that rapidly provides a high-precision solution of the initial phase under low excitation intensities (28). The resolution of the system is 105nm in X and Y and produces images at 300X, ideal for clear discernment of structures within prokaryotic cells (Supplementary Information Figure 3).

### Multiple SSB-foci are observed in exponentially growing cells

To visualize SSB under controlled expression levels, it was necessary to co-express an SSB-GFP fusion with wild type SSB. Under these conditions, chimeric SSB tetramers are produced (25). Tetramers containing up to two GFP fusion subunits retain wild type levels of activity (ssDNA and partner binding), while those containing three fusion subunits are reduced in their ability to bind interactome partners (25). To create the dual expression strain, MG1655 Δ*lacY::kan* was lysogenized using λDE3 to express T7 RNA polymerase under the control of the lac operon. Next, the region spanning the *ssb-gfp* fusion and chloramphenicol resistance gene from plasmid pCA24N-his-SSB-GFP was amplified in PCR in two separate reactions to facilitate recombineering into the non-essential, flagellar gene locus (29). The first added flanking sequences homologous to *flgA* (24.34 minutes) and the second added homology to *flgE* (24.39 minutes). These genes were selected as they are recombinogenic loci and are transcribed in opposite directions relative to the direction of DNA replication. Following recombineering and colony purification, GFP levels were assessed using a fluorimeter and found to be the same, within experimental error, at each locus (data not shown). Next, single colonies from strain 29003 (MG1655 Δ*lacY::kan, flgA:: ssb-gfp-cam*) and 29007 (MG1655 Δ*lacY::kan, flgE:: ssb-gfp-cam*) were grown separately to early log phase and imaged using wide-field epifluorescence to assess uniformity of expression of GFP. As both strains behaved similarly, only 29003 was used for subsequent studies.

First, cells were grown to early log phase in LB + 0.2% glucose, treated with mitomycin C (MMC) for 30 min, stained with the inner membrane dye FM4-64 and imaged using wide-field and at super-resolution using SIM. Representative pictures from each imaging modality are presented side-by-side to enable direct comparison (Fig. 1). In these images, the inner membrane is clearly indicated by red FM4-64 staining and within the cells 1-3 dominant SSB-GFP foci are observed at different positions. In the wide-field image, a heterogeneous smear of GFP signal is also evident in the cytosol in multiple cells (Fig. 1A, boxed cell, zoomed in panel C). In sharp contrast, in the super-resolution SIM image, the smear is resolved into five foci and an additional elongated structure 1.3μm long (Fig. 1D). Thus, in addition to the greater than 2-fold resolution enhancement in X and Y, the SIM enables visualization of fine detail not apparent in wide-field.

**Figure 1.**
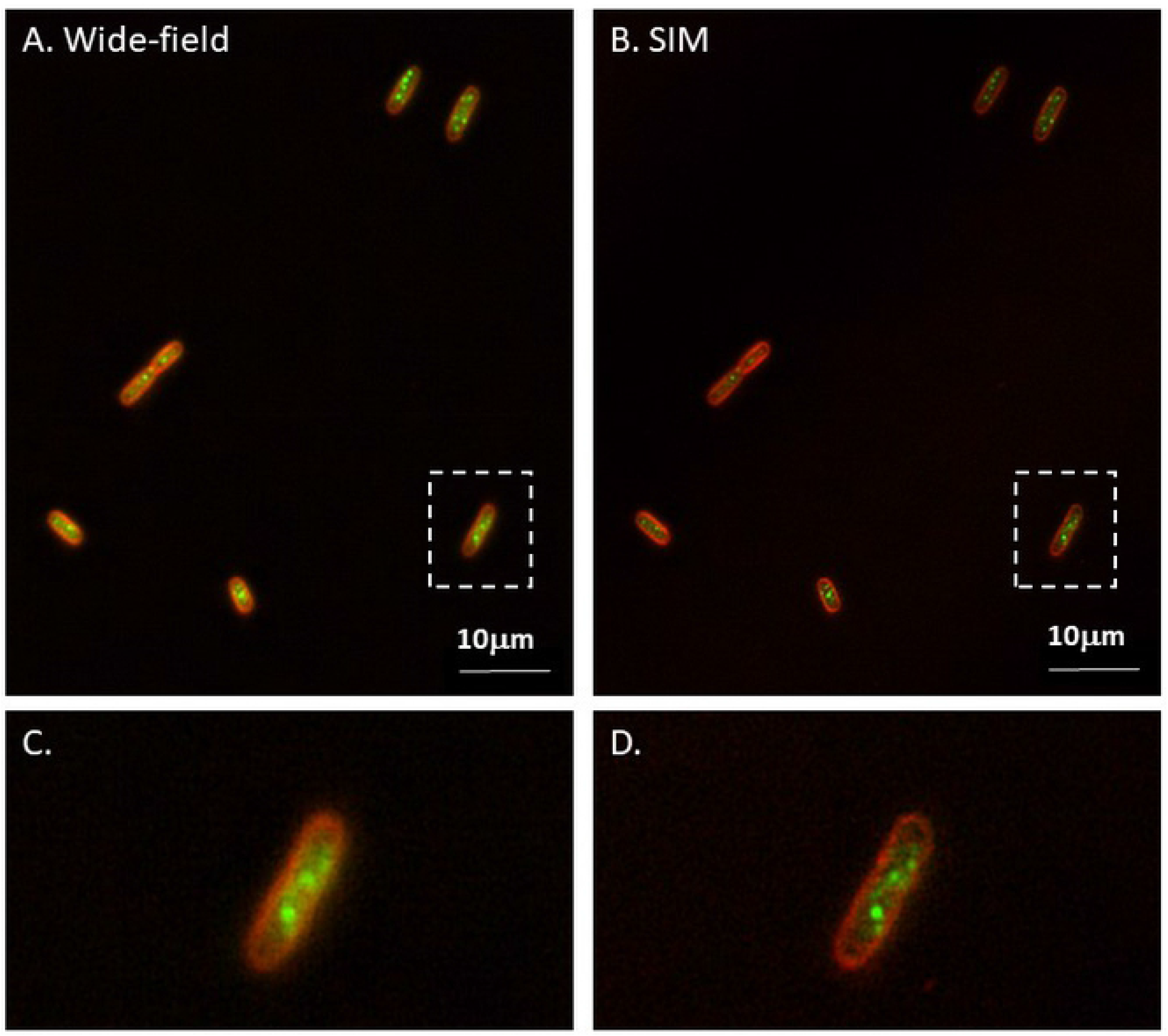
Super-resolution imaging reveals distinct SSB structures in exponentially growing cells exposed to DNA damage. To demonstrate the imaging capability of the SIM, cells were grown to mid-log phase in M63 media, treated with MMC, stained with FM4-64 and imaged using either wide-field (A) or structured illumination (B). For comparison, the same field of view is shown for each imaging modality. Panels C and D, zoomed images of the cells indicated by the boxed regions in panels A and D, respectively. For SIM images, raw data are shown. That is, no image processing was done, only the LUT was changed to red (FM4-64) or green (GFP).

Next, to visualize SSB localization in the absence of exogenous DNA damage, cells were separately grown to early log phase in either LB+0.2% glucose or M63+casamino acids +0.4% lactose and imaged. Glucose was used to lower the basal level of expression of the fusion in LB, while in M63, lactose was used as both a carbon source and low-level inducer of expression (data not shown). Using these growth conditions, dominant foci as well as a variety of additional foci and regions of enhanced GFP fluorescence in the cytosol were observed, independent of the growth medium used (Figure 2, compare panels A-C (LB) to panel D (M63)). Analysis of the boxed cell in Fig. 1C shows the presence of 3 dominant and 11 additional foci, many of which are located in proximity to the inner membrane. The row of five foci positioned near the inner membrane at the top of this cell are 200nm from the apex of FM4-64 fluorescence and are spaced 630 to 770nm apart (Supplementary information Fig. 4). As the distance between the apexes of GFP and FM4-64 signals in this region are 200nm, this means that in this location, SSB is not membrane-associated. In contrast, at the opposite membrane, the apexes of the FM4-64 and GFP signals are within 70nm of one another and some of the signal overlaps suggesting that SSB may be localized to the inner membrane in the absence of DNA damage as proposed previously (7).

**Figure 2.**
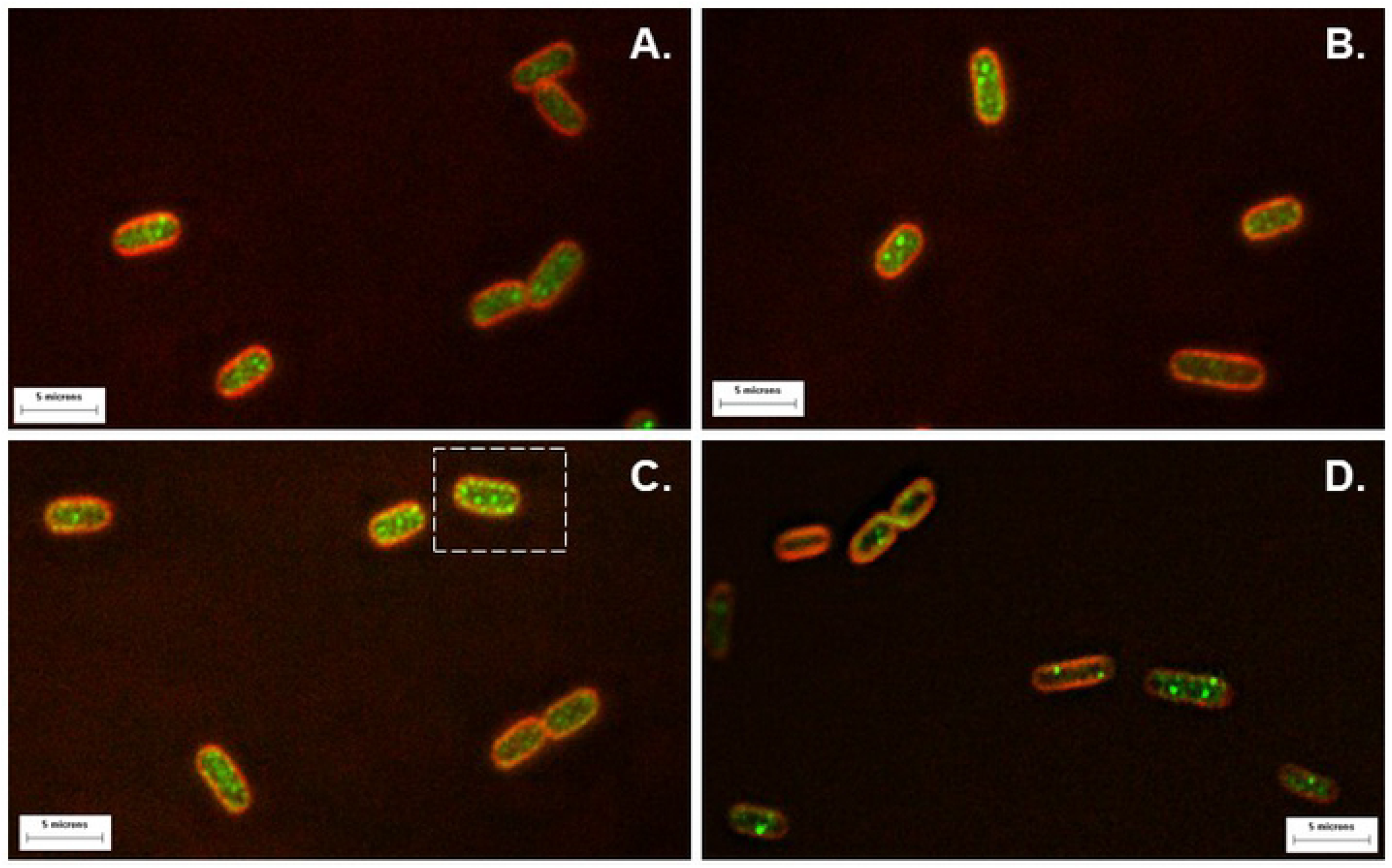
SSB is found at multiple locations in the absence of DNA damage. Representative super-resolution images of cells grown to mid-log phase in LB (panels A, B and C) or M63 (panel D). Structured illumination microscopy was done as described in Material and Methods. The resulting SIM images for FM4-64 and GFP were merged in Fiji to create the color images shown. Magnification is 300x, with relevant scale bars shown at the bottom of each image. The boxed cell in (C) was analyzed further (Supplementary Information Figure 3). For these images, raw data are shown. That is, no image processing was done, only the LUT was changed to red (FM4-64) or green (GFP).

### In the absence of DNA damage SSB localizes to the inner membrane

To assess membrane localization more precisely, the super-resolution images of cells grown in minimal media were analyzed. Results show that an average of 1.9±0.5 foci/cell were observed, with some cells lacking foci and others containing as many as five (not shown). In addition, in 74% of cells analyzed (31 total), the FM4-64 and GFP signals overlap over more than 50% of the inner membrane perimeter (Figure 3). In these overlap regions, the apexes of their signals are either within 46nm of each other or, their position is indistinguishable (Fig. 3C and F). Colocalization analysis of the boxed regions in Fig. 3G shows that in the overlap regions (boxes 1 and 2), the Pearson’s Correlation coefficients are 0.48 and 0.49, respectively. In contrast, within the center of cells, the correlation coefficient is either 5-fold lower (box 3) or there is no correlation between the FM4-64 and GFP signals (box 4). Therefore, in the absence of DNA damage a large fraction of SSB is localized to the inner membrane.

**Figure 3.**
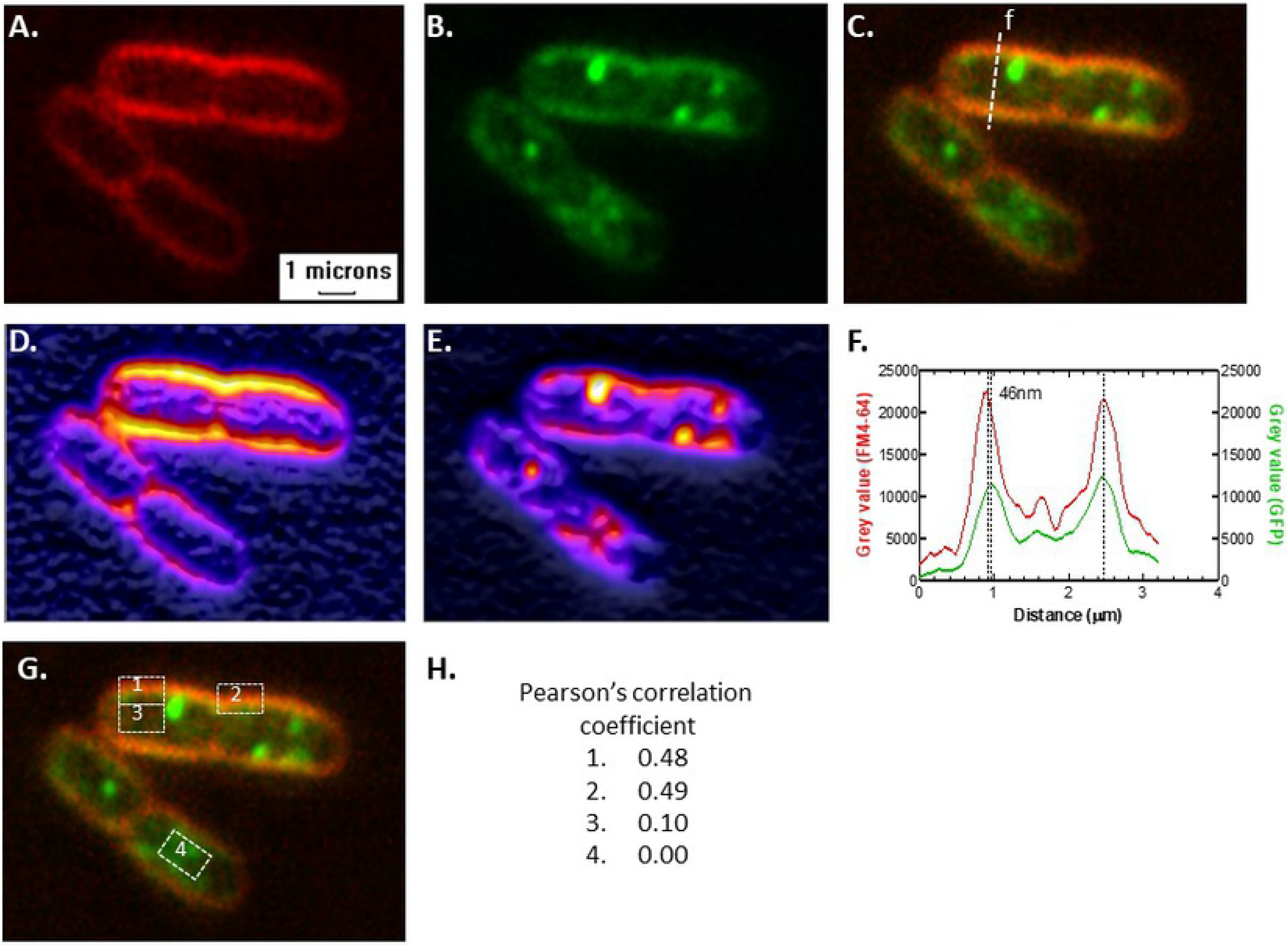
SSB is associated with the inner membrane in the absence of DNA damage. Representative SIM images of four cells from a mid-log culture grown in M63 are shown. A. FM4-64, B. GFP, C, Merge. D and E, 3D surface plots of the cells in panels A and B, respectively. F, Plot profile of line (f) indicated in panel C. The top of the line corresponds to 0μm distance. The dashed lines indicate the apexes of the FM4-64 and GFP signals. G and H, Colocalization analysis of the boxed regions indicated. For SIM images, raw data are shown. That is, no image processing was done, only the LUT was changed to red (FM4-64) or green (GFP).

### SSB binds phospholipids

The imaging data presented above suggest that SSB associates with the inner membrane in the absence of DNA damage. To determine if SSB binds to the phospholipid component of the membrane, a fluorescence-based assayed was used. Here, the intrinsic fluorescence of SSB, which is typically used to monitor binding to ssDNA, was instead used to monitor binding to liposomes, similar to that employed previously for RecA (27). In this assay, SSB fluorescence is monitored and aliquots of liposomes comprised of different compositions of phospholipids are added in separate reactions, and fluorescence measured. Liposome binding results in quenching of the intrinsic fluorescence of SSB (Figure 4A).

**Figure 4.**
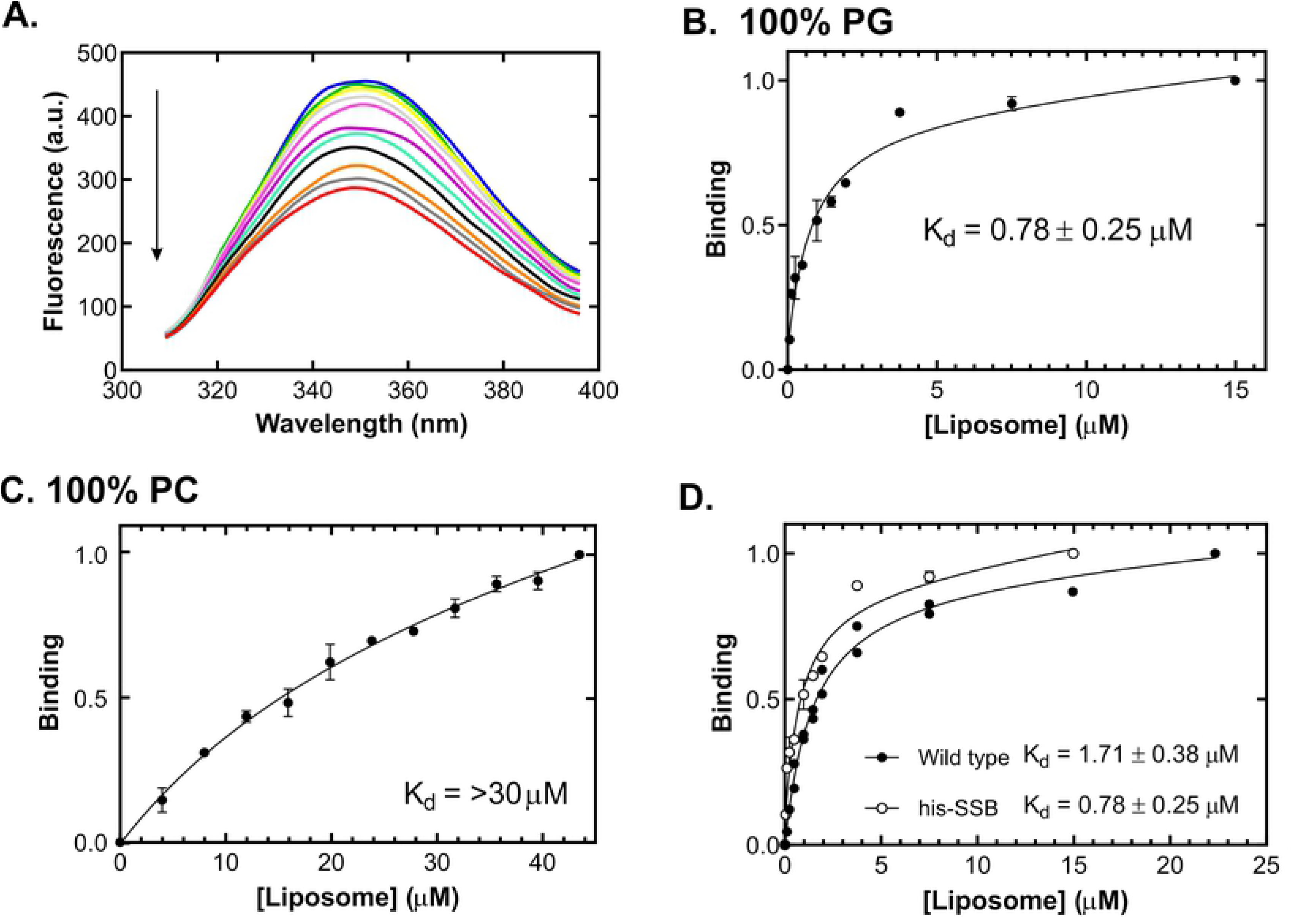
SSB binds to liposomes. A, Titration of SSB with liposomes comprised of 100% phosphatidylglycerol. A representative titration of 1μM SSB (monomer) with aliquots of 100% PG liposomes is shown. Assays were done at 37°C as described in the Materials and Methods. Each scan shown was done 8 minutes after the addition of liposomes to ensure that binding had occurred and a stable fluorescence signal was obtained. The arrow indicates increasing concentration of liposomes. B, SSB binds to 100% PG-containing liposomes. Analysis of data sets such as those shown in panel A is shown. Two separate assays done on separate days are shown. C, SSB does not bind appreciably to liposomes comprised of 100% phosphatidylcholine. D, Both wild type and his-SSB bind to 100% PG-containing liposomes. The data set for his-SSB is the same as that in (B). Data for all liposomes tested are summarized in Table I. Corrected data in each titration were fit to one-site specific binding equation: Y= (B_max_ × X)/(K_d_+X·) as described previously (27).

Results show that SSB binds to liposomes comprised of 100% phosphatidylglycerol (PG) with a K_d_ of 0.78 ± 0.25μM and very poorly to 100% phosphatidylcholine (PC) liposomes (K_d_ >30 μM; Fig. 4B and C and Table I). This makes sense as PG is a major component of the inner membrane, while PC is not present (30). Due to the instability of 100% cardiolipin (CL) liposomes due to the properties of CL, it was necessary to assess binding by mixing this phospholipid with the more stable PC. To enable direct comparison, binding to PG and PE were assessed as PC liposome mixtures as well, similar to what was done before for RecA (27). Addition of cardiolipin (CL), phosphatidylethanolamine (PE) or PG to phosphatidylcholine liposomes increased binding 38-to 86-fold resulting in the K_d_ values less than 1μM (Table I). As *in vivo* localization experiments were performed with strains containing both wild type SSB under the control of its own promoter and his-SSB-GFP expressed from an ectopic locus, binding to both Wt- and his-SSB was assessed. The results show that the affinity of each protein for liposomes comprised of 100% PG was within experimental error, the same (Fig. 4D).

These data show that SSB, like RecA and DnaA, binds to the anionic phospholipid components of the inner membrane (27, 31, 32). However, and unlike RecA, SSB also demonstrates an affinity for the zwitterionic phospholipid PE, as indicated by a K_d_ value of 0.56 ± 0.20 μM observed in the presence of 80:20 PC:PE liposomes (Table I). Further, the affinity of SSB for each of the preferred phospholipids is on average, 10-fold greater than that observed for RecA (27).

### Following DNA damage, the sub-cellular location of SSB changes

To determine whether membrane-localized SSB is mobilized in response to exogenous DNA damage, cells were grown to early log phase, treated with different concentrations of MMC for 30 minutes, stained with FM4-64 and imaged at super resolution. Results show that in the presence of 0.2 μg/ml MMC, cells contain a greater number of foci than in undamaged cells and that the majority of the GFP signal is present within these foci or associated with the genome in some fashion. Furthermore, the locations of SSB are distal from the inner membrane (Fig. 5A). In the presence of 1.2 μg/ml MMC, similar results were obtained although fewer cells survive the damage (Fig. 5B). Importantly, the level of SSB associated with the inner membrane decreases significantly following DNA damage to low levels that are not discernible as distinct peaks when quantified (Fig 6, panels B-D). In contrast, in undamaged cells, the GFP signal is observed a clear peak at the membrane (Fig. 3F). Some SSB still remains membrane-localized following MMC treatment however, but this was observed in only 9% of damaged cells analyzed (64 total). This is 8-fold lower than what was observed in undamaged cells where membrane-localization was observed in 74% of the cells analyzed. In these experiments, it is not possible to visualize the genome as the microscope used is a prototype with only one illumination laser.

**Figure 5.**
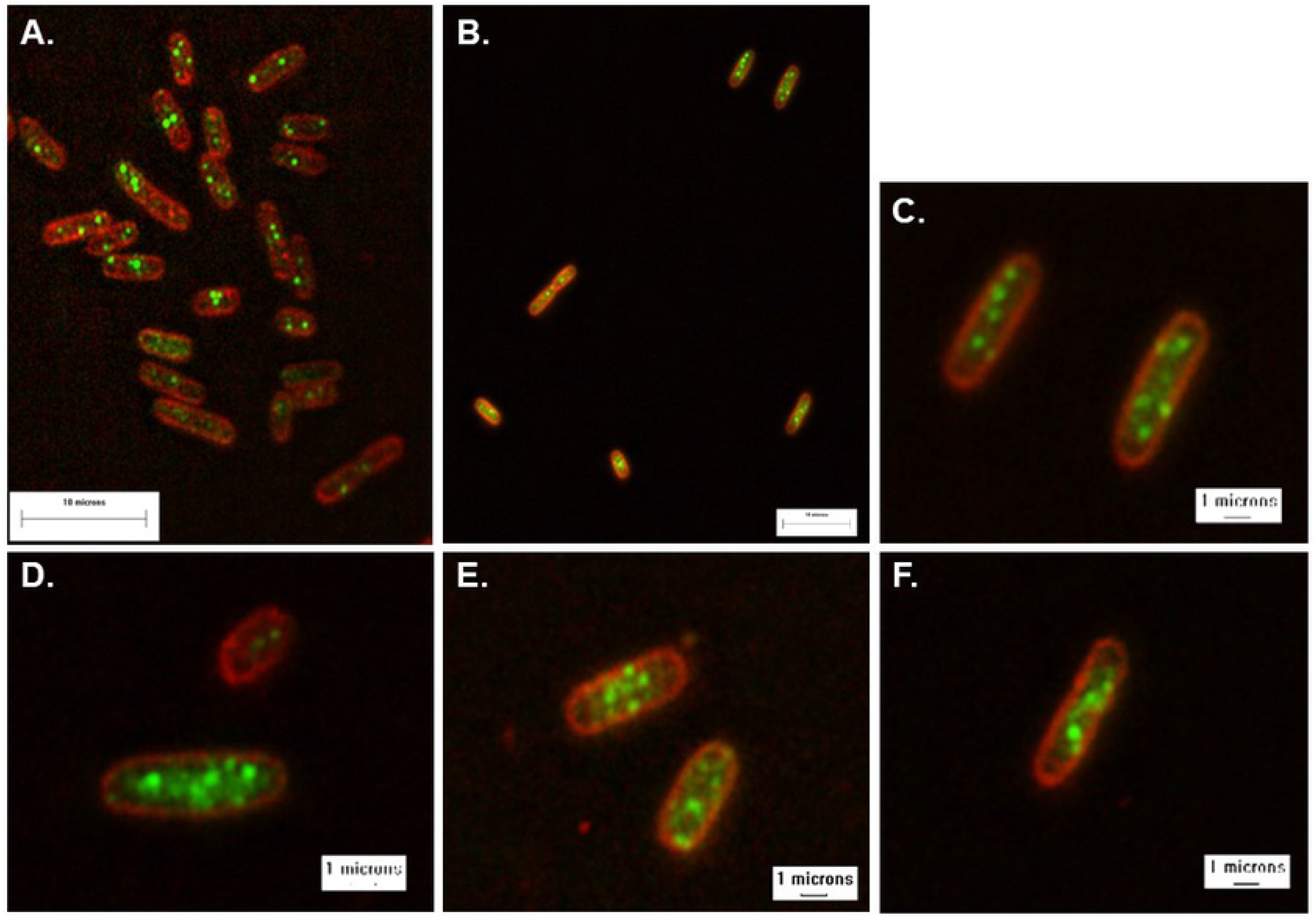
Treatment with mitomycin c induces changes in the sub-cellular localization of SSB. Cells were grown to mid-log phase in M63 media, treated with MMC for 30 minutes, stained with FM4-64 and imaged. The merged images are shown. A, representative cells from a culture treated with 0.2 μg/ml of MMC. B, representative cells from a culture treated with 1.2 μg/ml of MMC. Panels C-F, zoomed images of cells with damaged genomes. In these images, raw data are shown. That is, no image processing was done, only the LUT was changed to red (FM4-64) or green (GFP).

**Figure 6.**
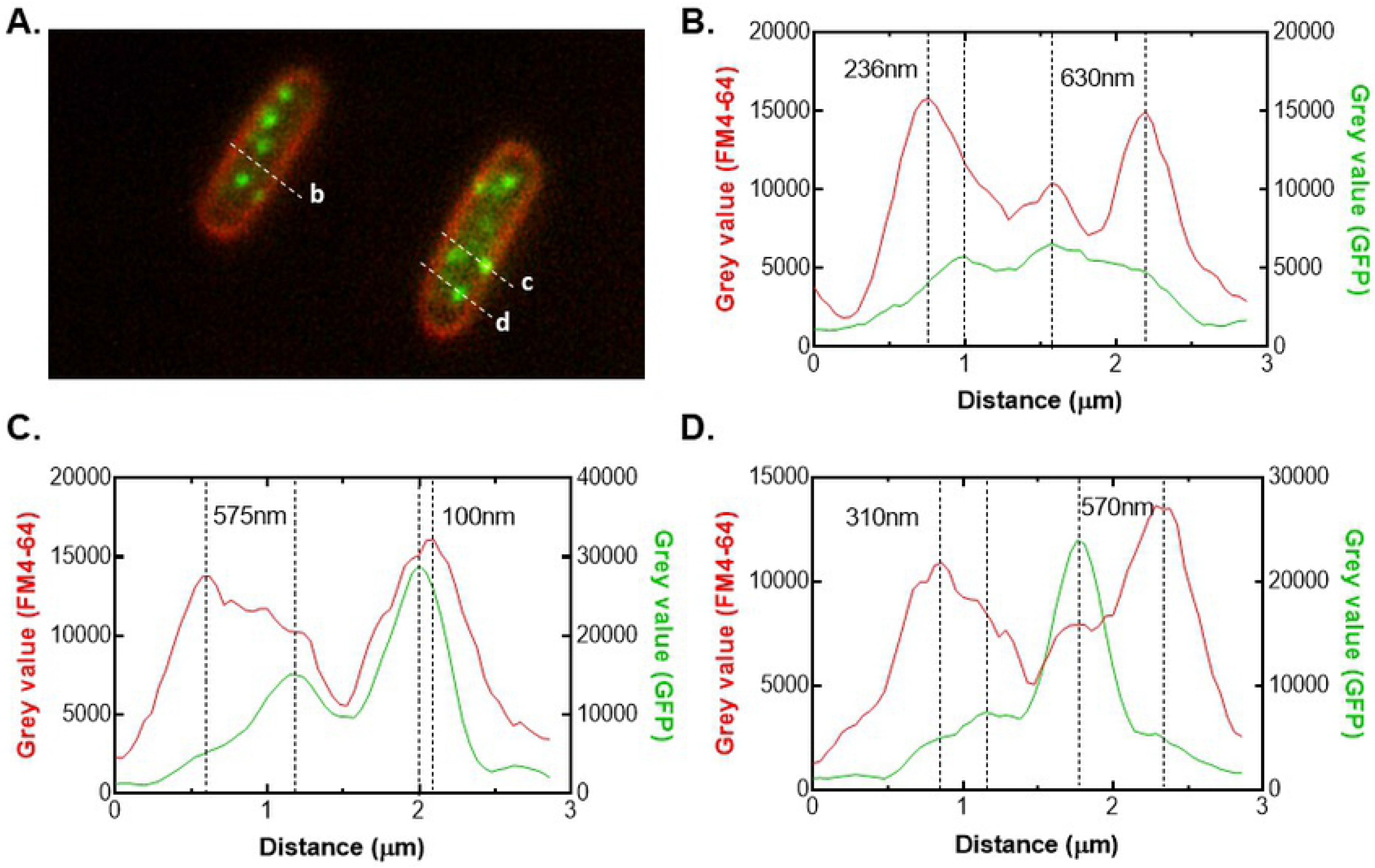
SSB is no longer associated with the inner membrane following DNA damage. The analysis of two representative cells from a culture exposed to 1.2 μg/ml of MMC is shown. The cells were taken from the top right of the image in Figure 4B. The intensity profile of the lines a, b and c are presented in panels B-D, respectively. The left end of each line corresponds to 0μm distance. The values between the vertical dashed lines in each plot indicate the measured distances between the apexes of the FM4-64 and GFP signals.

Next, a time course of exposure to 1.2 μg/ml MMC was performed. Here cells were again grown to mid-log phase, MMC added and 1ml aliquots of cells removed from the culture at various times. Immediately following their removal, aliquots were subjected to centrifugation, pellets resuspended in fresh LB, re-centrifuged and resuspended in 10mM MgSO_4_ and prepared for imaging.

Analysis of images of time courses reveals that at the earliest time point, that is 5 minutes after treatment, the level of membrane-associated SSB is significantly reduced and cells appear identical to those in Figures 4 and 5 which were captured 30 minutes after MMC treatment (data not shown). Attempts to determine how rapidly SSB returns to the membrane once MMC is removed, have proved challenging. In these experiments, cells were grown to mid-log phase, treated with MMC for 30 minutes, subjected to centrifugation to remove MMC, resuspended in fresh LB and grown for various time points (15; 30; 45; 60 and 120 minutes). By 15 minutes, membrane-localized SSB is present but this is only observed in less than 3% of cells examined (total 60). This value rises to 5-8% at 30-45 minutes, depending on the experiment (data not shown). By 60 minutes following MMC removal, cells are in stationary phase and the experiments were terminated. In those cells in which membrane-localized SSB was observed, the number of foci ranged from 1-3 per cell suggesting that repair had taken place and cells had recovered from the damage event (data not shown). In contrast, in the remaining cells, 5 or more foci were present in addition to various SSB structures indicating that damage had not yet been completed.

## Discussion

The primary conclusion from this study is that in the absence of DNA damage, a significant fraction of SSB is membrane-localized where it binds to the major phospholipids of the inner membrane. We propose that this corresponds to the storage form of SSB with perhaps as many as 1,900 tetramers, poised for action. Following DNA damage, SSB is rapidly mobilized so that within 5 minutes, it is no longer associated with the inner membrane but instead becomes associated with the genome enabling repair processes to occur.

The association of SSB with the inner membrane in our view, makes sense. It is known that the basal level of SSB in cells grown in LB is 8,232 molecules per cell or 2,058 tetramers (19). Using a site size of 40 nucleotides of ssDNA occluded per tetramer, 82,320 nts can in principle be bound by all of the available SSB (25). However, a typical DNA replication fork is thought to have 1,000 nts of ssDNA exposed suggesting approximately 25 SSB tetramers will be found per fork (20). In undamaged cells we see on average, 2-4 foci per cell. If these correlate with sites of DNA replication as expected, 100 SSB tetramers are bound to sites of replication leaving 1,958 free. The results herein show directly that the storage site of a large fraction of this free SSB is at the inner membrane. Membrane-association is not unique to SSB, having already been demonstrated for DnaA, RecA and UmuC (27, 33–35).

If free SSB is membrane-localized, how is binding mediated? It is well known that a key element of the SSB protein is the OB-fold (36). This fold is responsible for binding both ssDNA and the linker domain of adjacent SSB tetramers thereby enabling cooperative interactions (8, 37). Importantly the OB-fold is structurally similar to Src homology 3 (SH3) domains found in eukaryotes (11). Recently it has become clear that in addition to protein binding, SH3 domains also bind lipids (38, 39). The *in vitro* data presented herein show that SSB binds with high affinity to the three major inner membrane phospholipids PE, PG and CL (Figure 4 and Table I) (30, 40). Therefore, it is conceivable that like SH3 domains, the OB-fold of SSB is responsible for mediating phospholipid binding thereby enabling localization to the inner membrane. The low μM K_d_ values observed for the binding of SSB to the major inner membrane phospholipids likely ensures that the large pool of free SSB is not deleterious to the genome. In addition, tight binding is likely not problematic for SSB function as it binds to ssDNA with even greater affinity with K_d_ values as low as 3nM (41). Consequently, we suggest that lipid binding may be competitive with ssDNA binding, providing an additional level of control over SSB function similar to that proposed for c-Src regulation in eukaryotes (39).

The results herein show that within 5 minutes of being exposed to MMC, SSB has disengaged from the inner membrane and becomes associated with the genome. This association is visualized as a dramatic increase in the number of foci per cell as well as the presence of both clearly discernable structures (Fig. 5F) and clouds of GFP signal spread over the genome (Fig 5D). These clouds may correspond with multiple sites of DNA damage requiring the presence of SSB and/or the protein diffusing to sites where its activity is required. The large structures on the other hand may correlate with regions of the genome having suffered significant DNA damage requiring extensive repair. Due to its importance in multiple DNA transactions it is not surprising that SSB rapidly disengages from the membrane in response to exogenous damage. In contrast, UmuC is sequestered at the inner membrane until it is activated by RecA* nucleoprotein filament-mediated cleavage of UmuD to UmuD’ (35).

Once damage has been repaired, SSB once again localizes to the inner membrane. Due to technical reasons, it was only possible to visualize the return phase in a limited number of cells. Here, the number of foci per cell was 2 and the GFP signal was found associated with the membrane similar to what is observed in undamaged cells (Figure 2). Therefore, the results herein show that SSB cycles on and off the membrane in response to the damage status of the genome. This is distinct to what was proposed for RecA where the inner membrane was shown to have a role in nucleating RecA filament bundles which is important for SOS function (27). Instead, we propose that membrane–localization sequesters excess SSB away from the genome thereby minimizing excessive strand separation and/or spurious melting of duplex DNA that otherwise might be lethal to the cell (42–44).

In this study we show that excess SSB is localized to the inner membrane, where it is sequestered until needed. Due to its critical role in DNA metabolism it disengages within 5 minutes to facilitate repair processes where it binds to ssDNA rapidly and with high affinity and returns once repair has taken place. As SSB is known to bind to the repair helicases RecG and PriA *in vivo* in the absence of DNA (7), it is conceivable that following damage the SSB-RecG/PriA complexes disengage from the inner membrane and SSB targets the helicases to sites of DNA replication fork stalling, resulting in rescue. Studies are underway to determine whether this does in fact occur.

## Acknowledgements

Work in the Bianco laboratory is supported by NIH grant GM10056 to PRB. Work in China was supported by the Natural Science Foundation of China (NSFC) grant 61522511 to ML. Work in the Nguyen laboratory is supported by NIH grant EB023262 to JN.

